# siQ-ChIP: A reverse-engineered quantitative framework for ChIP-sequencing

**DOI:** 10.1101/672220

**Authors:** Bradley M Dickson, Rochelle L Tiedemann, Alison A Chomiak, Robert M Vaughan, Evan M Cornett, Scott B Rothbart

## Abstract

Chromatin immunoprecipitation followed by next-generation sequencing (ChIP-seq) is a key technique for mapping the distribution and relative abundance of histone posttranslational modifications (PTMs) and chromatin-associated factors across genomes. There is a perceived challenge regarding the ability to quantitatively plot ChIP-seq data, and as such, approaches making use of exogenous additives, or “spike-ins” have recently been developed. Relying on the fact that the IP step of ChIP-seq is a competitive binding reaction, we present a quantitative framework for ChIP-seq analysis that circumvents the need to modify standard sample preparation pipelines with spike-in reagents. We also introduce a visualization technique that, when paired with our formal developments, produces a much more rich characterization of sequencing data.

## Introduction

The “immunoprecipitation blues” have been hard to shake for groups that are reliant on the chromatin immunoprecipitation followed by next-generation sequencing (ChIP-seq) method for mapping the distribution of histone posttranslational modifications (PTMs) and transcription factors across the genome[1]. These blues result from concerns regarding antibody behavior (specificity and IP efficiency) and from a perceived challenge regarding the ability to quantitatively plot ChIP-seq results. Regarding the latter, a host of methods have been introduced in efforts to add a meaningful “y-axis” to ChIP-seq datasets[2, 3, 4, 5, 6, 7], calls to arms have been issued[6, 8, 9, 10], and additional ChIP methods presenting newer solutions are introduced regularly[11]. Herein, we apply physics-based mathematical modeling to derive a quantitative framework for ChIP. In this attempt to shake the blues, we demonstrate that ChIP-seq is (and always has been) quantitative without the need to modify standardized sample preparation pipelines[12] with “spike-in” reagents.

Fundamentally, the IP step of ChIP is a competitive binding reaction. As such, this step can be described through biophysical models used to define binding constants and explain competition reactions.[13] Upon developing such a model, a perfectly quantitative and spike-in free approach emerges where the only new requirements for the experimentalist are to carefully track the common variables of biophysical measures (*i.e.,* concentrations, volumes, masses, and buffer compositions) used in each step of commonly adopted ChIP-seq protocols[12]. The only real assumption is that the nature of physical interactions between an antibody and on- and off-target epitopes is the same everywhere on Earth. Under this assumption, any two ChIP results will fall on the antibody binding isotherm provided that reaction conditions are maintained across the experiments so that essentially only the distribution of epitope varies.

ChIP-seq appears non-quantitative because a fixed mass of DNA is always taken to library preparation regardless of how much DNA was captured by IP. This practice ensures that the number of obtained reads matches the number of reads that were requested. Obtained reads, represented as the green line in Figure 1A, are invariant to experimental conditions. The number of total reads held in the IP can be much larger (or smaller) than requested. Our approach explicitly makes use of the “down-sampling” (or “up-sampling”) of material in order to relate sequenced reads to the total possible reads. This is the bridge between the fundamental laws of binding that play out in the IP and the invariant sequenced read depth. Undoubtedly, the amount of material obtained by IP follows the sigmoidal curve shown in Figure 1A, a complete expression for which is obtained below. This is the antibody binding isotherm and, even though it cannot characterize any single microscopic binding constant, it can be used to characterize the general mass-balance that occurs in bulk. A schematic of a ChIP-seq protocol is shown in Figure 1B along with several factors that are determined during the experiment.

**Figure 1.**
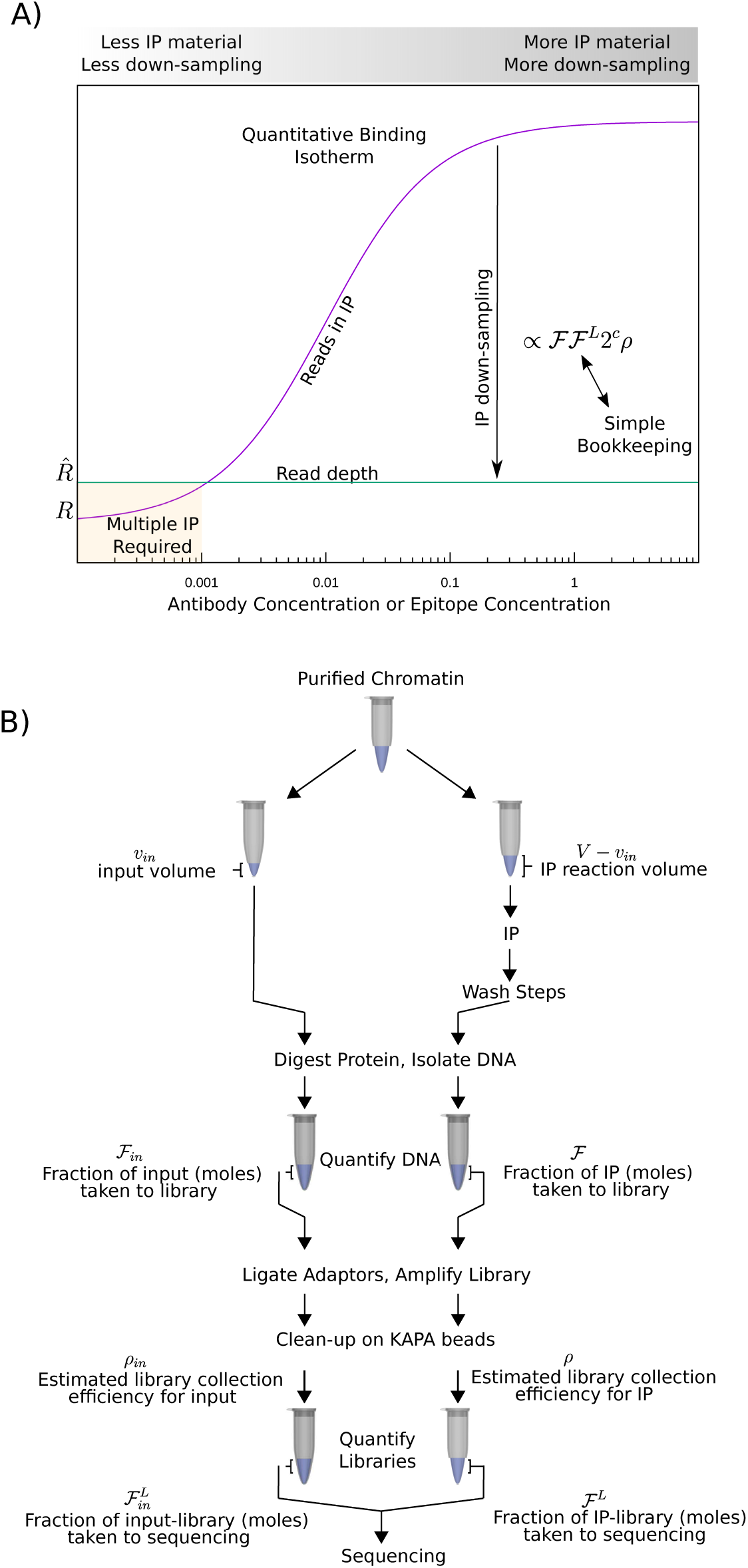
Key concept. A) The sequenced reads (or fragments) is a subset of the total possible. Down-sampling to meet the requested depth causes loss of quantitation. Restoration of quantitation only requires some bookkeeping during ChIP-seq practice. B) Schematic of bookkeeping.

The specifics of binding can be refined from this global perspective by viewing the sequencing results as a record of labels for all particles retained by the IP. The identity of a particle is vaguely, but well enough, characterized by three attributes: (1) genomic coordinates, (2) fragment (or “read”) length and (3) estimated capture efficiency. Once the binding isotherm is projected down onto the sequencing results, the familiar and microscopic notions of binding constants emerge and tightness of binding can be plotted as a function of genomic location. As this high resolution view of binding processes is not compatible with conventional genome browser visualization, we also represent ChIP-seq data in alternative ways. In example comparative analysis of ChIP-seq datasets presented below, this new visual representation is shown to produce actionable indications of on- or off-target binding that are lost in traditional genome-browser views.

Formally, the model defines how the number of reads in an IP depend on the concentrations of antibody, on- and off-target species concentrations and all binding constants between antibody and on- and off-target species. Even though these parameters are unknown in practice, this approach provides significant leverage for understanding ChIP-seq outcomes. Model predictions were validated against the empirical behavior of antibody specificity and global normalization, as reported by semi-synthetic spike-in nucleosomes (which are a proxy for all exogenous spike-in normalizers).

Applicability of our approach, called <u>s</u>ans-spike-in method for <u>q</u>uantitative ChIP-seq (siQ-ChIP), only requires that any ChIP-seq experiments that are to be compared should be performed in the same reaction volume, same input volume, same amount of initial chromatin, and the same amount of antibody — a common yet implicit demand of all (before-IP) normalization schemes. The results described below are generalizable to cases where different numbers of cells must be used to harvest chromatin and also to cases where multiple IP reactions must be combined to reach workable chromatin amounts for library preparation, maintaining the ability to quantify results in these cases. For completeness, explicit mention of combining multiple IPs is made in closing.

Below we use the words “read” and “fragment” interchangeably. siQ-ChIP is specifically designed for paired-end sequencing, so mixing read and fragment is a tolerable abuse of notation as long as the reader keeps this in mind.

## Results and Discussion

As a benchmark system for developing a quantitative ChIP-seq formalism, we conducted ChIP-seq for lysine 27 tri-methylation on histone H3 (H3K27me3). Chromatin was harvested from HCT116 cells treated with vehicle (DMSO) or EPZ6438, a potent inhibitor of the catalytic subunit of the Polycomb Repressive Complex 2, EZH2.[14] Complete experimental details are found in Supporting Information. EZH2 is the primary enzyme known to catalyze H3K27me3 in mammalian cells. Thus, EPZ6438-treated cells present limited target epitope relative to vehicle-treated cells. For IP, we selected a commonly used H3K27me3 rabbit monoclonal antibody (CST C36B11; lot 9733S(14)), shown to be highly specific by proxy of histone peptide array, publicly available at www.histoneantibodies.com.[8]

The SNAP-ChIP[4, 9] method was used for these experiments as a comparative, spike-in method for quantification. Small amounts of semi-synthetic labeled, or bar-coded, mononucleosomes, unmodified or bearing different orders (mono-, di-, or tri-methylation) of H3 and H4 methylation states (H3K4, H3K9, H3K27, H3K36 and H4K20) were spiked in with intact chromatin prior to micrococcal nuclease (Mnase) digestion. These exogenous spike-ins are intended to provide a global scaling factor so that ChIP-seq data can be normalized and compared. Additionally, these spikeins provide an estimate of antibody binding specificity that affords some metric of cross-reaction between antibody and nucleosome species. These nucleosomes are referred to as SNAP nucleosomes below.

This experiment led to two observations which motivated this work: (*i*) counterintuitive observations of antibody specificity reported by spike-in nucleosomes and (*ii*) ambiguously interpretable spike-in normalized sequencing tracks. In brief, antibody specificity was observed to improve when performing ChIP with chromatin from EPZ6438-treated cells, contradictorily suggesting that the antibody is more specific for target when it is in excess of the target. Additionally, the normalized track could equally well be interpreted as either evidence for the formation of new H3K27me3 peaks in EPZ6438-treated cells or as having no signal depending entirely on the biases of the user. In fact, as we explain below, all of the current globally-normalized schemes[3, 4, 6] lead to a similar compression of signal and potentiate the same ambiguous interpretation.

The situation is summarized in Figure 2. Three approaches to quantify ChIP-seq data are shown. Line graphs depict what is visualized in a genome browser for each approach, and heat maps depict the three components of particle identity that underlie the browser line: (1) genomic coordinate, (2) fragment length, (3) IP:input, histone modification density (HMD)[4, 9], or estimated capture efficiency *ê*(*x, L*). The information embedded in these heat maps is extremely valuable for data interpretation but are never constructed and are incompatible with conventional genome browsers. While fragment start site and length are the same for all normalization methods, the third attribute, giving the color scale to the heat maps and the height of the line graph, is unique to each approach. Only siQ-ChIP returns an actual estimated capture efficiency yet this notion is at the heart of each estimate.

**Figure 2.**
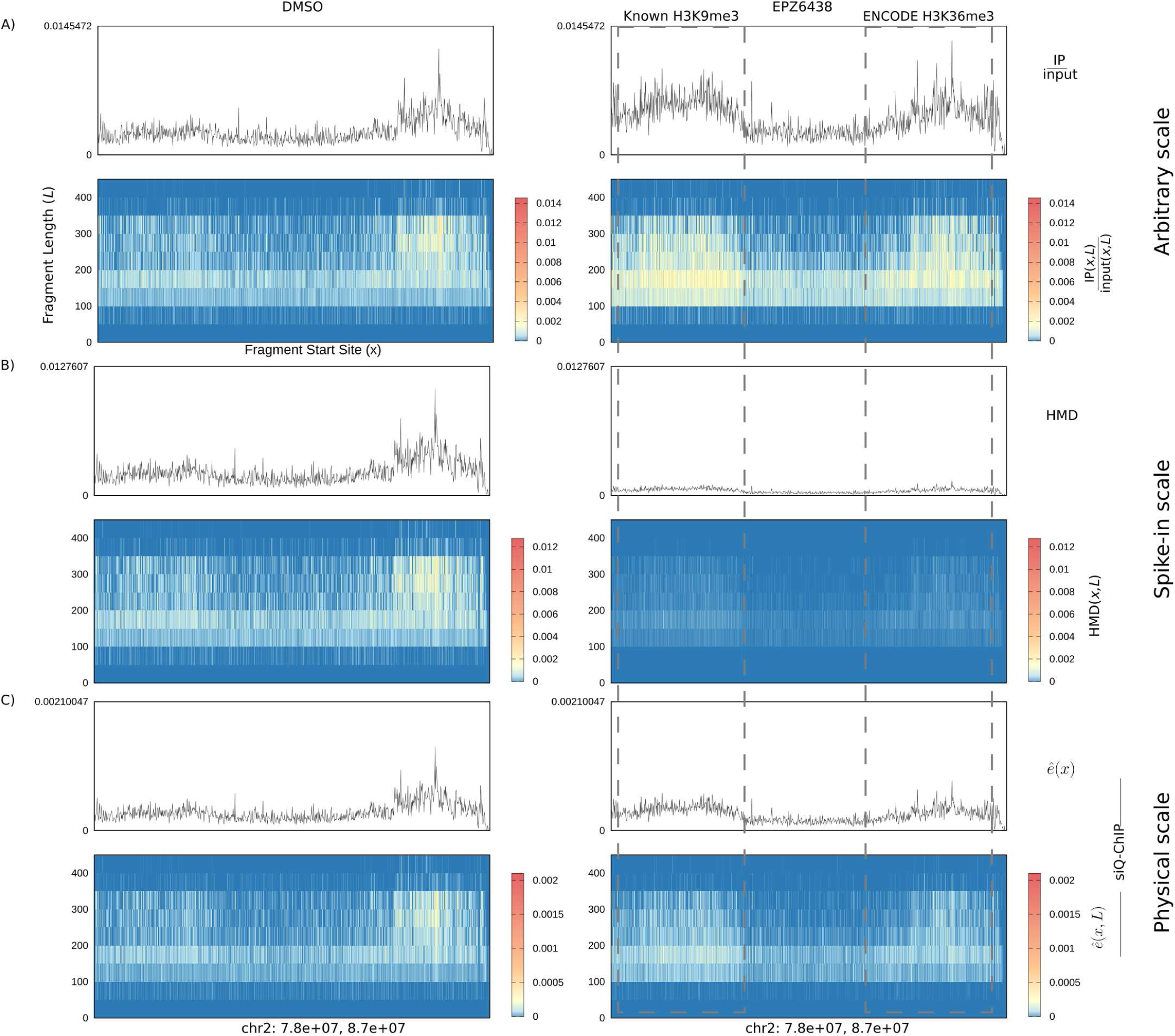
A question of scale. IP:input normalized only to number of mapped reads, Histone modification density (HMD), and siQ-ChIP. Single lines show genome-browser style data while heat maps show histograms of fragment as a function of start position (x-axis), and length *L*, where color gives scaled estimates of IP efficiency.

Notably, in the case of DMSO treatment, all three methods show strong agreement in shape with only the scale of the heat map and line graph being different. Any peaks attributed to H3K27me3 in one method would also be seen in the other methods, and any peak-to-peak comparison within an experiment would be invariant to the approach used. The data differ only by absolute “normalization,” which is simply multiplication by a constant.

Likewise for EPZ6438 treatment, the methods strongly agree in pattern, but not in scale, and here the discrepancy is exaggerated. Thus, for experiments performed under the same reaction conditions (*i.e.,* same reaction volume, antibody concentration, chromatin mass), as was the case for these two experiments, the issue of quantifying Chip-seq data is simply a matter of determining scale.

Which of these scales is the “correct” one? Relating the IP to a binding model first defines the context in which we can interpret correctness of scale and forces adherence to a protocol that isolates changes in scale to relative changes in concentrations of chromatin epitopes. Simultaneously, it also produces a much deeper understanding of the ChIP-seq output. For example, we show below that the two peaks in EPZ6438 results, which are obtained by IP with an H3K27me3 antibody, are correlated with ChIP-seq of H3K9me3 (left peak) and H3K36me3 (right peak). The siQ-ChIP (sans-spike-in method for <u>q</u>uantitative ChIP-seq) heat map scale holds enough information to imply the off-target nature of these peaks, which is how we found these peaks in the first place. Our model for ChIP-seq also shows that the off-target nature of these peaks leads to the signal compression displayed by HMD (“histone modification density”) in Figure 2B. The extent of compression is so great that one would presumably conclude the HMD track is empty. Given the monetary expense of ChIP-seq, ignoring data is tantamount to throwing money away. Not to mention, the HMD track is in direct conflict with the fact that Figure 2A suggests a large amount of material was captured in the IP. The ability to deconvolute the nature of peaks and extract maximal information from results should be a top priority of any analysis. Our method allows extraction of greater information content by avoiding compression and allowing the scale of IP efficiency at each genomic location to be an independent observable.

The next section establishes the basic model for binding and shows how it connects to mapped reads (or paired-end fragments).

### Binding relations for IP and input

Sequencing of the EPZ6438-treated IP produced a total of 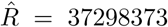 mapped reads. These reads were generated by sequencing 20 femtomoles of library, where the total library mass was 856 femtomoles. Setting ℱ^*L*^ = 20*/*856 = 0.02336 for the fraction of the library that was sequenced, the total number of reads that the full library would generate upon sequencing is 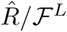, or 1.5 billion for this experiment. This is the first step in estimating the number of effective reads that were captured in the raw IP, a number important to the connection between sequenced reads and the binding phenomena producing the epitope-enriched IP material. The library was amplified with *c* = 11 cycles of PCR, so this estimate of the total reads must be reduced by the appropriate number of amplifications, lowering the estimate of total reads to its pre-amplification value 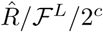.^[1]^ Furthermore, the library was captured on KAPA Pure beads producing an additional material loss, *ρ. ρ* is the ratio of captured library concentration to the expected library concentration.^[2]^ This coefficient compensates for losses due to bead capture, washing, and, to some extent, for global deviations from the perfect 2^*c*^ amplification. The estimated number of reads becomes 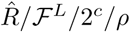. The observed read count 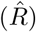 has been scaled up by each known source of material loss.

Similarly, if the IP produced 24.2 ng of material and 10 ng were used to produce library then, where ℱ = 0.413 is the fraction of IP material carried into the library, then 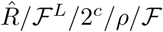 reads would be generated by sequencing all of the DNA collected by IP.

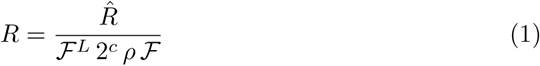

Thus for one of the experimental replicates presented herein, an estimated 226 million reads could be extracted from the IP material. Equation (1) can be re-arranged to relate the observed reads, 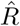 to the effective reads held by the IP

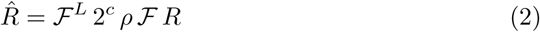

The hat symbol indicates that an instrument was used to convert a raw quantity *R* to a measured quantity 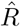.

As shown in Figure 1, equation (2) maps the theoretical reads in IP to the empirical constant 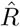. Any single ChIP-seq IP is one point on the isotherm of Figure 1 as antibody and chromatin are rarely titrated. (See Reference [15] for the exception and notice that it conforms to Figure 1.)

The key hypothesis motivating this work is that if *R* can be derived from the phenomenological description of binding, common to all biophysical methods for determining binding constants, then a quantitative framework for ChIP-seq can be developed. Below, a standard description of binding equilibria is used to define *R* as a function of antibody concentration, epitope binding constant, off-target binding constants, and relative concentrations of all epitope species that interact with the antibody (*i.e.,* non-zero binding constants). The following development culminates with the extension of the model to genomic data, where heterogeneity plays a fundamental role. The equilibrium limits of the ChIP binding reaction are characterized in Supporting Information (Figure S2).

For *N* species that interact with the antibody, the conservation of mass requires that the total antibody concentration (*AB*^*t*^) equal the sum of free antibody (*AB*^*f*^) and antibody bound to all the interacting species (*S*_*i*_)

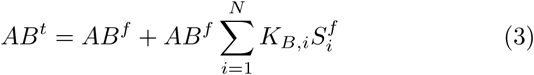

The following constraint also applies

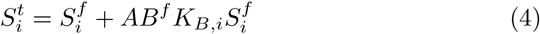

for all *N* species, where 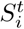 and 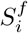 are the total and free concentrations of species *i*, respectively. The free concentrations of each component of the system can be found as the simultaneous solution to these equations if the total concentrations and binding constants are known. In ChIP-seq experiments, nearly none of this information is known, but this model provides relationships that will be useful for ChIP-seq analysis nonetheless.

For each of the 1 ≤ *i* ≤ *N* species, the free concentration is

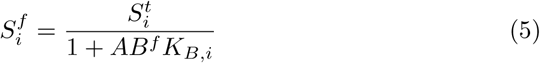

The concentration of species *i* that is bound by antibody is

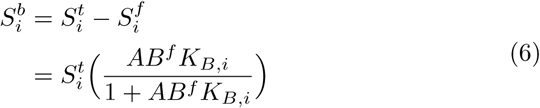

Equation (6) shows us that the amount of species *i* that is captured by the antibody depends on the amount of free antibody, which by Equation (3), is a function of all the binding constants and concentrations of all of the off-target species. Thus, the amount of bound target epitope depends on the relative amounts of all epitopes that the antibody can bind. Note that Equation (6) is the classical logistic function, sometimes called the Langmuir isotherm, associated with binding: When *AB*^*f*^ = *K*_*d*_, exactly half of 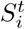 is bound. (*K*_*d*_ = 1*/K*_*B*_ is the dissociation constant.) This function is shown in Figure 1. Thus, we interpret binding constants as macroscopic avidity constants consistent with the formalities of mono- or poly-valent interactions. See for example the development of the enhancement factor *β* by Mammen, Choi and Whitesides.[13]

The number of moles of species *i* that were captured can be found by multiplying the concentration 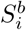 by the volume of the IP. Let the IP volume be *V* - *v*_*in*_ where *v*_*in*_ is a small volume removed prior to introduction of antibody. The volume *v*_*in*_ is the input volume. The number of moles captured by IP can be turned into a number of particles using Avogodro’s number *N*_*A*_.

If we assume that every particle of 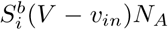 can generate one read, then Equation (1) can be used to evaluate the number of reads that 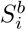 will generate according to our previous estimates of experimental losses

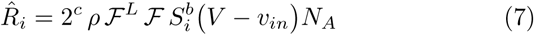

The total reads from IP, 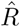, is given by adding up the reads 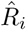 from each species captured by the antibody.

Likewise, the total reads in IP are

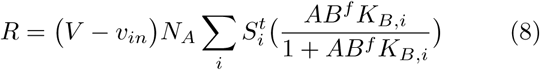

Through Equation (6), the predicted number of reads for each species is rigorously connected to the full spectrum of antibody binding constants, the antibody concentration and the relative concentrations of the various epitopes. Equation (2) can now be cast with explicit connection to the parameters governing the antibody binding reaction

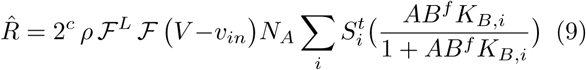

A prediction for the reads generated by *i* in input can be obtained similarly

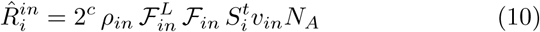

Table 1 summarizes all the coefficients that must be tracked throughout the ChIP-seq experiment. These are the coefficients that set the natural, quantitative scale in ChIP-seq experiments. A tabular summary of all measurements required to build these coefficients is given in the Supporting Information (Table S1).

**Table 1.** Table of symbols required for quantification. See Supporting Information for an expanded table of all required measurements.

| Symbol | Description |
| --- | --- |
| $\mathcal{F}$ | Fraction of IP (moles) taken to library |
| $\mathcal{F}^L$ | Fraction of IP-library (moles) taken to sequencing |
| $\rho$ | Estimated library collection efficiency for IP |
| $\mathcal{F}_{in}$ | Fraction of input (moles) taken to library |
| $\mathcal{F}_{in}^L$ | Fraction of input-library (moles) taken to sequencing |
| $\rho_{in}$ | Estimated library collection efficiency for input |
| $v_{in}$ | input volume |
| $V - v_{in}$ | IP reaction volume |

Equations (7) and (10) afford a formal expression for the efficiency of the IP for individual species

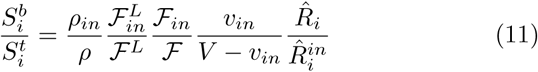

This expression for efficiency connects all the reads of species *i* to the concentration of *i* that was bound by the antibody, 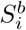, and the total unreacted concentration of *i*, 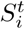. Using Equation (6) for 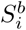 in Equation (11) reveals a connection to the binding constant *K*_*B,i*_

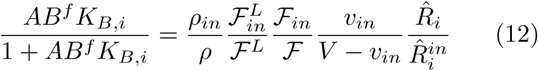

With the following definition of capture efficiency

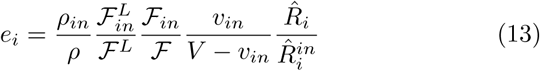

equation (12) can be rearranged to produce 

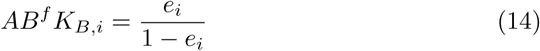

The solution to the system of equations given by (3) and (4) produces only a single value for *AB*^*f*^, the concentration of unbound antibody. Thus, the ratio of binding constants within an experiment can be estimated by

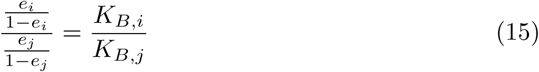

because factors of *AB*^*f*^ will cancel. Equation (15) is especially interesting because for carefully crafted experimental conditions it should take the same value at any antibody concentration. This expression is the only such invariant in the literature and should be considered when characterizing antibody quality and composition. A closing remark toward this aim is made in the Conclusion section.

Figure 3 summarizes the results of simulating two competing species in two practical contexts: A wild-type context (DMSO-treated) and a target epitope-depleted context (EPZ6438-treated) where the concentration of target (H3K27me3) is drastically reduced. The simulation also included trace amounts of spike-in nucleosomes that present on- or off-target epitopes. The binding constants for on- and off-target species were set to provide a 100-fold preference for target epitope. All simulation details are found in the Figure Legend.

**Figure 3.**
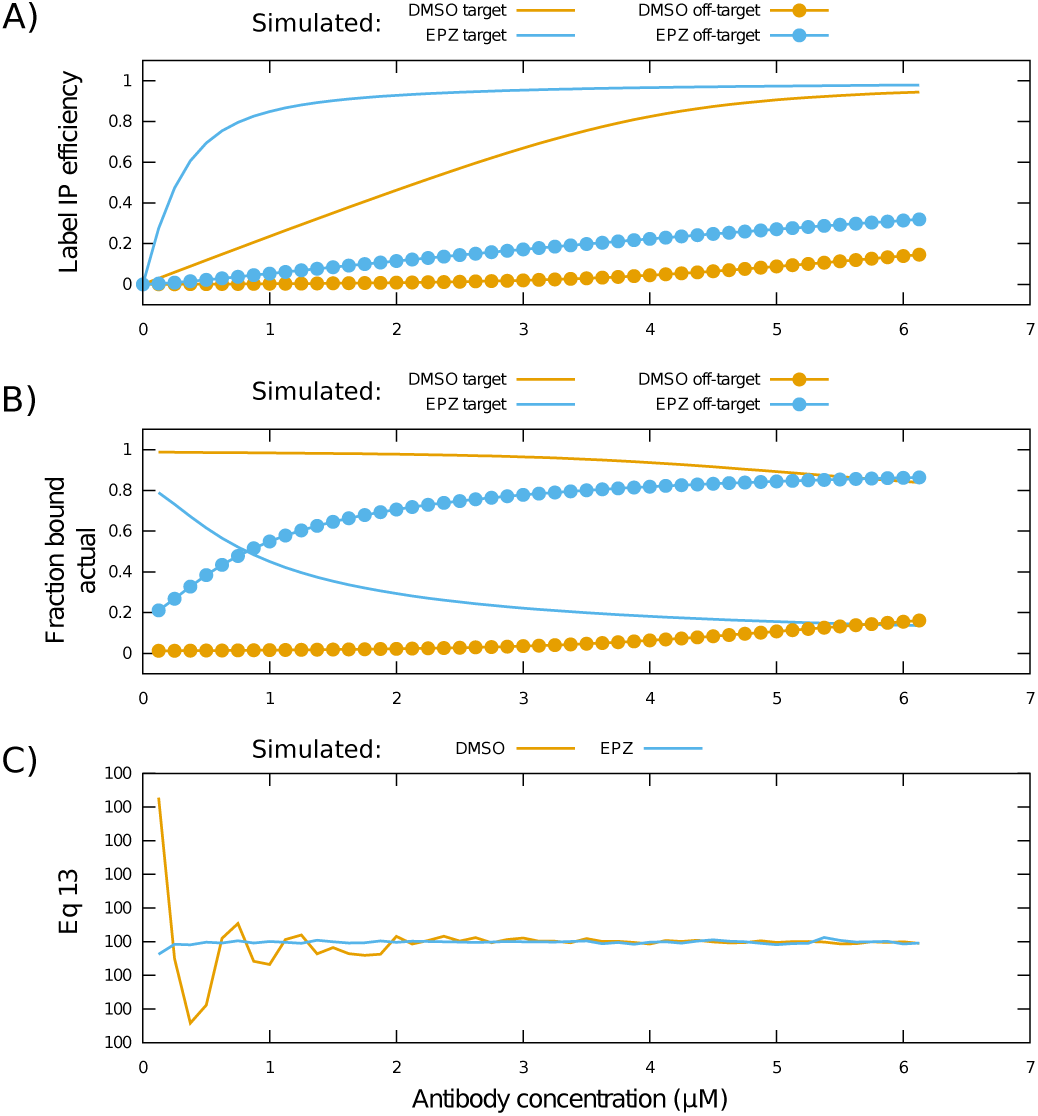
Model simulation. Simulated data for two species model. (A) IP efficiency for labeled spike-ins. (B) Actual fractional composition of bound unlabeled species. (C) Estimated ratio of on-target to off-target *K*_*B*_, variation is in parts per trillion. Actual ratio was 100, *K*_*B,*1_ = 11, *K*_*B,*2_ = 0.11. In the reaction volume labeled spike-ins were 0.06 *µ*M, unlabeled off-target epitope was 5 *µ*M, target epitope was 4 *µ*M in mock DMSO and 0.2 *µ*M in mock EPZ6438 treatment. No attempt was made to exactly fit experimental outcomes below.

The simulation predicts that capture efficiency of the labeled on-target spike-in will improve drastically when unlabeled chromatin is deprived of target epitope (Figure 3A). Additionally, the off-target labeled spike-in capture efficiency is not predicted to change to the same extent.

Figure 3B shows that when target epitope is depleted, the majority composition of bound particles shifts toward off-target species as the antibody concentration is increased. For low antibody concentrations, where the antibody concentration does not exceed target concentration, the bound particles are primarily on-target. Consistent with intuition, the simulation predicts that off-target binding is increased when antibody is in excess of target epitope. Figures 3A-B therefore predict that the capture efficiency for labeled target spike-ins can be significantly higher than for labeled off-target spike-ins. Yet, if the chromatin is primarily composed of off-target epitopes (as in the target-depleted case) then most of the captured chromatin will be off-target. This is not indicated by the spike-ins. Consistent with their use in wet-bench experimentation, the labeled spike-ins constitute a small fraction of the total particles in these simulations. Therefore, if label capture efficiency is 10% then we expect roughly 10% of the corresponding unlabeled epitope is also captured. The mismatch in interpretation comes because 10% of spike-in is far fewer nucleosomes than 10% of unlabeled nucleosomes, *i.e.,* most nucleosomes are unlabeled. In the context of epitope depletion, a 10% capture of 5 *µ*M off-target species produces a larger particle count than a 95 % capture of a 0.2 *µ*M on-target species. Simulations predict that even when the antibody is “specific,” one can expect to capture off-target epitope in an IP depending on the relative concentrations of all species in the IP. Moreover, the spike-in nucleosomes do not report on the number of bound unlabeled nucleosomes which allows for misinterpretation. Figure 3C shows that Equation (15) produces a context-independent estimate of the ratio of binding constants. We will discuss how Equation (15) could be used to empirically evaluate relative binding constants for antibody cross-reaction in the Conclusions.

Equations (11) through (15) are the first key results of this work. These expressions relate the total number of reads of a species to the respective binding constants and provide significant leverage for comparing ChIP-seq data as shown below. Some technical considerations are required yet to cast these results onto genomic coordinates, as these expressions so far apply only to the total number of reads generated by species with finite binding constant to the antibody. In a typical ChIP-seq experiment one would not likely be able to estimate 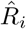 or 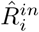, because it is impossible to group reads by species. Instead, one would have access to reads as a function of genomic location. We next relate our binding model to outcomes for the semi-synthetic labeled spike-in nucleosomes, which can be sorted by species, as a first step toward generalizing our model to genomic coordinates.

### Extension to labeled semi-synthetic nucleosomes

Consider projecting equation (11) onto labeled exogenous semi-synthetic nucleosomes. For any nucleosomal species that has a barcoded counterpart, the total species can be broken into the sum of labeled and unlabeled parts 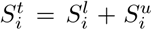 where *l* and *u* denote labeled and unlabeled. The *observable fraction* of 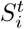 is 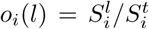, which is the labeled (*i.e.,* observable) fraction of the species. Using barcoded spike-ins to report on antibody-chromatin interactions assumes that the interaction with epitopes from labeled and unlabeled (endogenous chromatin) species are the same. Under this assumption, if a total of 𝒩 nucleosomes of species *i* are bound by the antibody in the IP one intuitively expects roughly *o*_*i*_(*l*) of those nucleosomes to be labeled. The concentration of bound label can also be found by multiplying Equation (6) by *o*_*i*_(*l*), 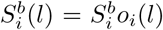, allowing reads of species *i* to be projected onto the labeled subset of the species

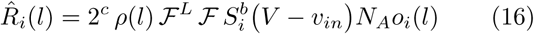

For the labeled input, we have simply

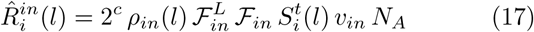

because the concentration of labeled spike-in is independent of the unlabeled concentrations. By protocol, the same amount of labeled spike-in is always added to chromatin, so the input cannot report on anything but the labeled concentration which is always the same. The amount of label in the IP, however, is connected to the concentrations of unlabeled chromatin through the competitive binding reaction that is summarized by equations (3) and (4). These estimates of labeled reads can be substituted into Equation (11) to find the IP efficiency of the labeled fraction of species *i*

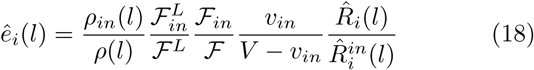

The hat on *ê*_*i*_(*l*) distinguishes the measured efficiency for the label *l* from the theoretical efficiency *e*_*i*_ for species *i*. We use *ρ*(*l*) to indicate the library losses specifically for the spike-ins. In practice there is no good way to estimate this number and one does not need spike-ins to determine scale so below we willingly take the assumption that *ρ*(*l*) = *ρ*_*in*_(*l*).

Under antibody-saturating conditions, 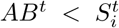, the reads captured for label *l* on target species 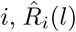, can be decreased by increasing the unlabeled concentration 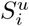 through the action of *o*_*i*_(*l*) in Equation (16). In this limit of concentrations the amount of captured particles 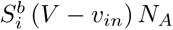 will not change (since all of the antibody is already bound), yet the amount of bound label 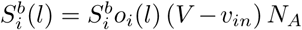 is diminished as 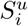 is increased simply because it is more likely that the antibody encounters unlabeled target. This is intuitive as it reflects the consumption of antibody by increased unlabeled chromatin levels. It is precisely this behavior that disallows the appearance of *o*_*i*_(*l*) in the estimate of input reads 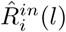 because the input reads do not depend on 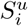.

The model, as we have defined *ê*_*i*_(*l*), predicts that the capture efficiency for the on-target H3K27me3 spike-in will approach unity when EPZ6438 inhibitor is used to deplete H3K27me3 levels. The observability of the target label increases with this treatment because the unlabeled concentration of epitope 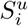 decreases, thus 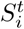 approaches 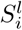 and 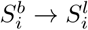. This prediction was experimentally validated with the SNAP-ChIP spike-ins, where we measured a 25% capture efficiency for on-target semi-synthetic nucleosomes spiked in DMSO-treated chromatin versus a 95% capture efficiency when spiked in EPZ6438-treated chromatin.

To further validate the model with spike-in nucleosomes, additional model-based predictions were tested by experiment. The model affords a predictive framework for several common empirical metrics and how they respond to epitope depletion. Consider the ratio of efficiency for an off-target species *j* and an on-target species *i*

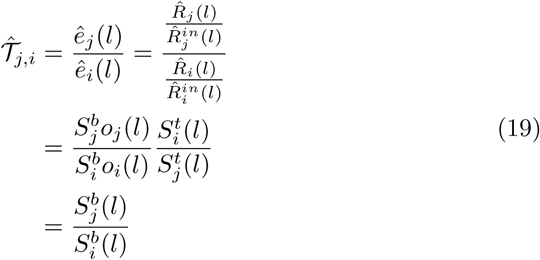

Note that the experimental prefactors from Equation (18) cancel from numerator and denominator here because 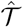 is a ratio of efficiencies. Recall that by protocol 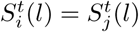.[4, 9]

The estimator 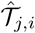 is used to qualify an antibody as “good,” meaning specific, or “bad,” meaning non-specific, as follows: For any off-target (*j*) this estimator is expected to be small for a high quality antibody and comparatively large for a low quality antibody.[4, 9] We used Equation (18) in Equation (19) to write the specificity estimate 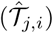 in two alternate representations that reveal dependence on species concentrations and observability, allowing predictions to be made regarding the behavior of 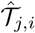 in the context of epitope depletion. When EPZ6438 is used to deplete H3K27me3, the quantity 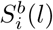 is expected to approach saturation. This is shown in Figure 3A (solid blue line). In cells that do not have depleted epitope, 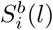 is pre-dicted to be under saturation (solid gold line). Thus, if the value of 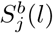 is roughly independent of the target epitope concentration, as would be expected for weakly interacting off-target species, 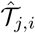 is expected to decrease upon epitope depletion by a factor proportional to the increase in the quantity 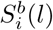. That is, an amount proportional to the amount of target label bound. Thus, the model predicts a compression of the 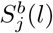 signal reported by 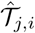 for any off-target *j*. Reducing 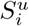 or increasing 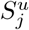 are predicted to artificially improve observed specificity 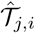 without requiring an actual decrease in off-target binding 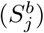, even though no physical attributes of the antibody have changed. Interestingly, if cellular chromatin presented equal amounts of all PTMs, this signal compression would not be observed, because the factors *o*_*i*_(*l*) and *o*_*j*_(*l*) would cancel from 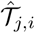. Of course, PTMs are distributed very differently, and those distributions change under experimental perturbation.

The other common metric of specificity (in the context of labeled nucleosomes) is

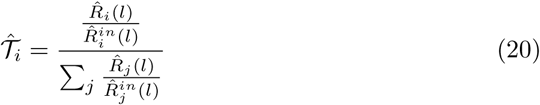

This metric estimates the fraction of on-target signal (or reads) out of the total signal, which includes any off-target reads. This metric is predicted to behave in parallel with 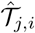 for the same reasons just discussed.

Figure 4 shows the results of Equation (19) for the same antibody in ChIP-seq from either DMSO- or EPZ6438-treated cells. The treatment with EPZ6438 is known to inhibit EZH2 and lead to a global reduction of H3K27me3, constituting widespread epitope depletion. As discussed above, the loss of target epitope reduces 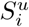, where *i* is the target species, leading to a predicted improvement in observed specificity due to signal compression. Indeed, Figure 4 shows improved specificity for the antibody upon epitope depletion. This shows that the measured specificity is a function of the unlabeled chromatin and does not report on the extent of inherent context independent cross-reactivity of the antibody (*i.e.,* binding constants). Also, as predicted, the IP efficiency of the H3K27me3 SNAP nucleosome (Equation (18)) is 25% in the presence of DMSO-treated chromatin and 95% in the presence of EPZ6438-treated chromatin, showing that there is excess antibody in the case of EPZ6438 treatment. Intuitively, performing ChIP-seq in the presence of excess antibody should increase the likelihood of observing cross reaction between the antibody and off-target species. However, the profiled antibody specificity has improved when antibody is put in excess. The context-dependent variation in measured specificity, and the counterintuitive nature of the estimate, are the source of disparate observations previously reported between peptide array and solution-based assays where the relative amounts of all species are not controlled between assay platforms.[16, 9] This counterintuitive behavior of 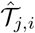 was predicted by the model.

**Figure 4.**
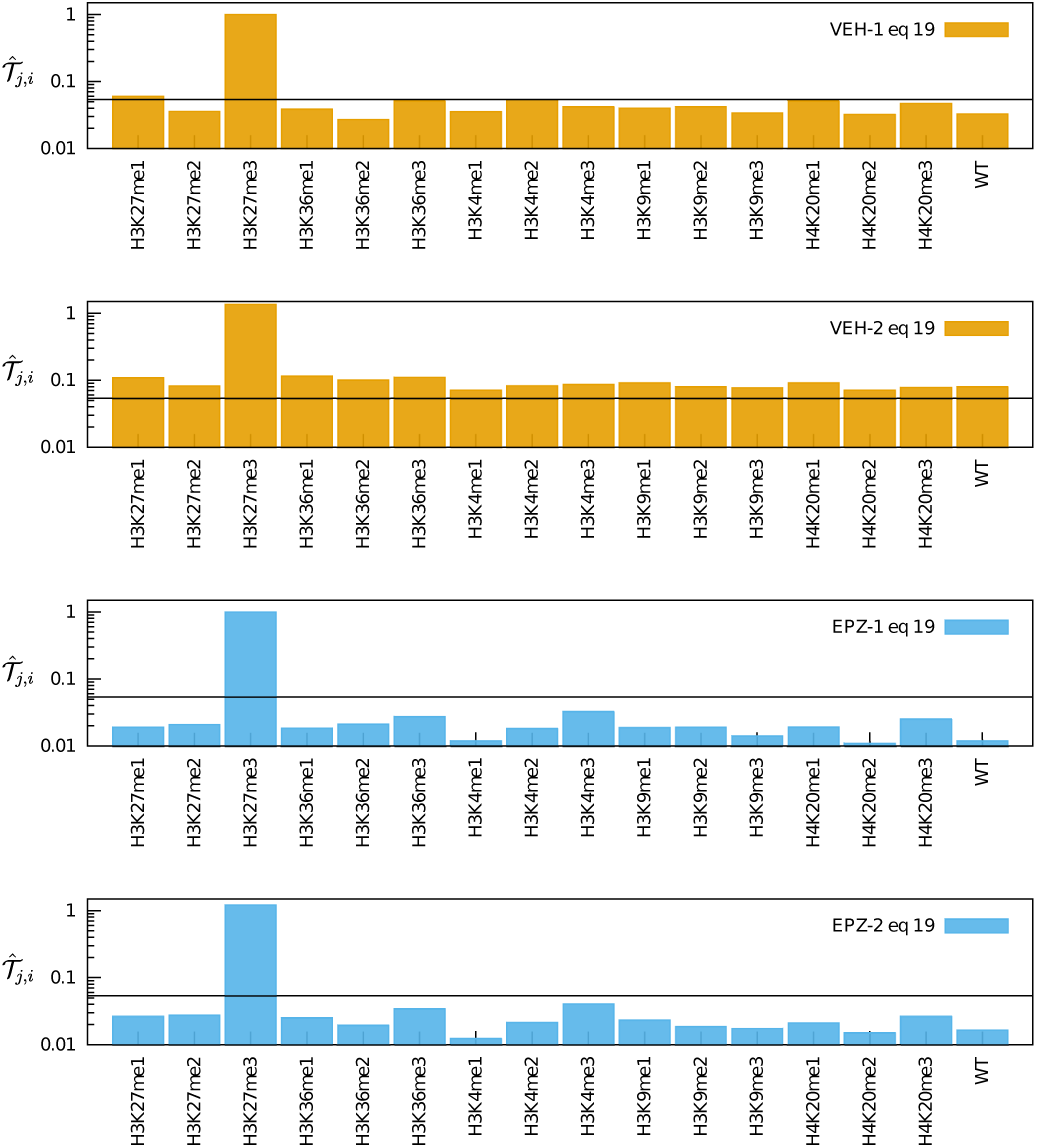
Observed selectivity 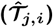. SNAP nucleosome selectivity profile in ChIP-seq from vehicle (DMSO) or EPZ6438 treated cells, two biological replicates. This is the so-called “% Target” specificity measure. Constant value of 0.05 shown in black as a visual reference. Data is average of SNAP pairs in one biological replicate. Full table in Supplemental Information. The target epitope, species *i*, is K27me3. Antibody: CST C36B11 lot 9733S(14)

Figure 5 shows the ratio of capture efficiency for each species with and without EPZ6438 treatment. This is the capture efficiency of each species compared to itself before and after EZH2 inhibition. In each replicate the capture of off-target species has increased in absolute quantities, as expected, with H3K27me3 depletion. However, this increase is masked in the specificity metrics of 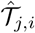 and 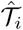.

**Figure 5.**
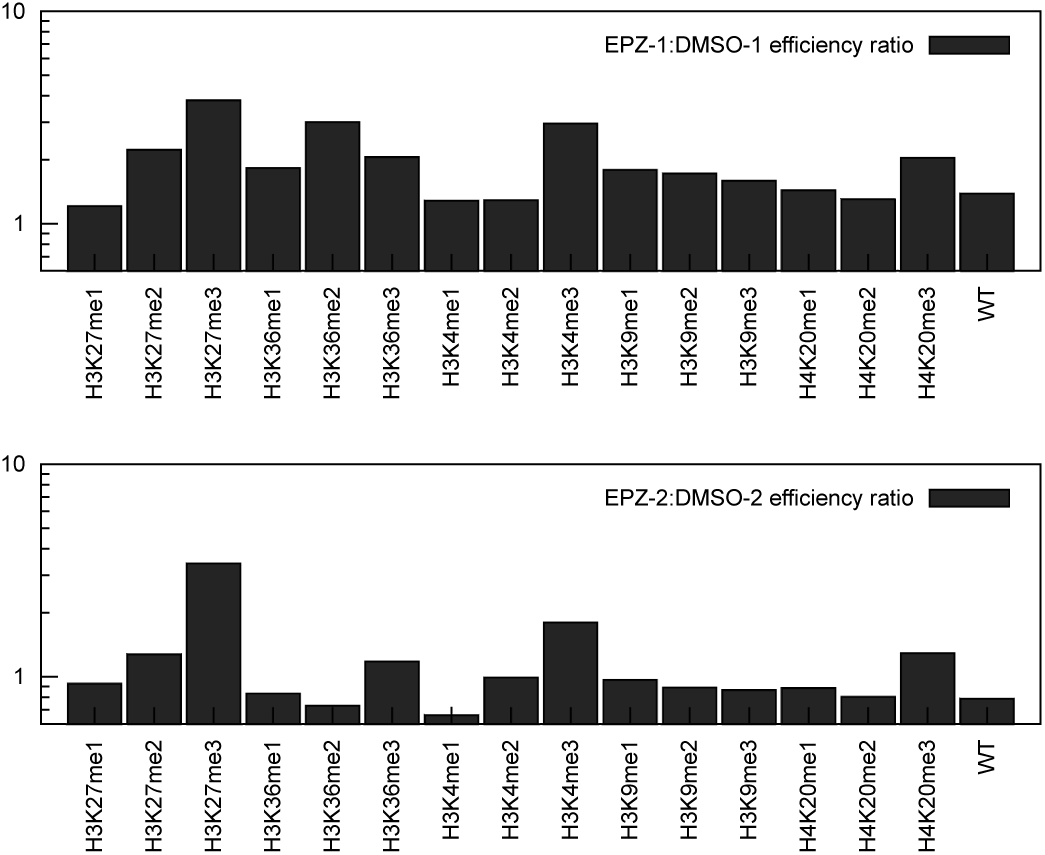
EPZ6438-dependent increase in semi-synthetic off-target capture. Ratio of efficiency, EPZ6438 to DMSO (vehicle), two biological replicates, for each spike-in species.

Additionally, note that Figure S4B of reference [9] shows that specificity given by Equation (20) can be improved from roughly 65% to 80% by adding unlabeled off-target epitope, consistent with the above model-based predictions.

The model has produced accurate predictions for labeled synthetic chromatin in two empirically tested quantities. Next, the model is extended to genomic coordinates.

### Extension to genomic coordinates

The set of cellular nucleosomes that constitute the particles of species *i* are scattered throughout the genome. Thus the reads of *i* are also scattered. Most importantly, reads at a particular genomic coordinate may not arise solely due to binding with a single epitope species. Due to the heterogeneity within a cell population, a mixture of PTMs could arise at a single genomic location, and all nucleosomes at a particular location may not be modified. This section investigates the extension of Equation (18) to genomic coordinates in light of heterogeneity.

Let a genomic region be denoted by *x*, where *x* specifies the position of a window of fixed base width. Now *x* can be thought of in the same way as the label *l* for exogenous nucleosomes. There is an observable fraction *o*_*i*_(*x*) for any species found at *x* where any of species *i* that is not at *x* is understood as unlabeled. The IP efficiency at *x* is only more complex than it was for synthetic labels because we must sum over all the species present at *x* to account for all the reads piled up at *x*. Firstly, the IP reads due to species *i* is

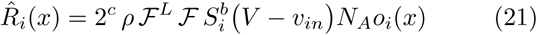

For input we follow all the above arguments to obtain

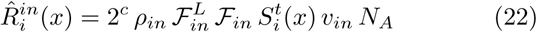

Again, input is not dependent on *o*_*i*_(*x*). Second, the total reads at *x* from IP is given as the sum 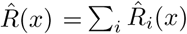. For input, 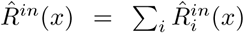. ChIP-Lseq experiments only track reads as a function of *x* and cannot exactly track which species generated the reads.

The reason reads are obtained at *x* in IP could be through interaction with on- or off-target, or a mixture of both, when population heterogeneity is considered. Thus, the only accessible estimate of IP efficiency is

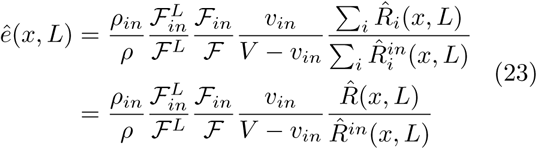

Two important details must be acknowledged here: First, we are making explicit use of paired-end sequencing which provides the length (*L*) of each fragment that is mapped to *x*. Second, in application of this expression, *x* is the interval within which a fragment starts. We find that *ê*(*x, L*) is only sensitive to the length of the interval *x* when the interval is taken too small. Basically the data become noisy, limiting the smallest interval width to around 100 base-pair for the data analyzed below. The three identifying attributes mentioned in the Introduction are *x, L*, and *ê*(*x, L*). Note that the total reads is now given by 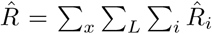, which relates everything back to Equation (2).

*ê*(*x, L*) can be transformed for plotting in a genome browser as the cumulative efficiency on an interval *y*

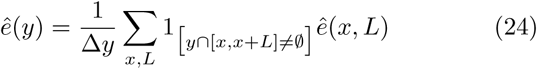

where the indicator function 1_[·]_ is unity if the interval *y* intersects the current fragment and is zero otherwise. The length of the interval *y* in base-pairs is given by Δ*y* so that *ê*(*y*) can be seen as the cumulative efficiency per base-pair. The cumulative efficiency (*ê*(*y*)) was shown in Figure 2. For visualization we also make an analogous projection of *ê*(*x, L*) onto *y*, which was shown in figure 2.

Before moving away from Equation (24), which is the basis of siQ-ChIP, we briefly make note of how our approach interacts with standard peak calling algorithms. There are some deep implications for peak calling, outlined in below, but it is worth noting that siQ-ChIP can be combined with standard peak callers. The widely used MACS peak caller[17], which calls peaks in part by the ratio of IP to input reads can be used here. If MACS is used to call peaks with a threshold *IP/input* > *θ* then the minimal peak height in siQ-ChIP is given by 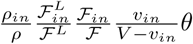. Any hidden Markov model peak caller would also be suitable. It is suggested to call peaks on the underlying data and then use those locations to examine/process *ê*(*x*). However, as discussed below, it is completely reasonable to plot the ratio of cumulative efficiency (Equation (24)) from two experiments that are being compared. This provides immediate access to regions of differential enrichment without using any peak callers.

### Application to ChIP-seq analysis

Figure 2 summarizes evaluation of *ê*(*x, L*) and *ê*(*x*) for a region of chromosome 2 and demonstrates how number of mapped reads and HMD normalizations compare. Looking back on Figure 2 the signal compression of HMD can be explained. The HMD normalization scheme is given in spirit by 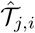 above, where species *j* is replaced with genomic reads at *x*,

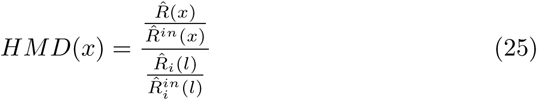

where *i* = H3K27me3 and *l* is the associated DNA barcode. The analysis of HMD is identical to that of 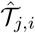, with one additional complexity stemming from cellular heterogeneity. In the case of HMD, signal compression can arise for both off-target binding, as it does in 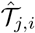, and also for on-target binding in regions of high heterogeneity. Because any exogenous chromatin can be built as a specific combination of synthetic nucleosomes, these results for HMD imply that all chromatin spike-ins will demonstrate comparable compression. The above implies that HMD does not explicitly express anything concrete about the “modification density” because the nature of compression is undetermined in practice. While it is generally appreciated that number-of-mapped-reads normalization is insufficient for capturing proper ChIP-seq scale, it is now clear that exogenous spike-ins also produce erroneous scaling. The denominator in Equation (25) was 1.14 for DMSO-treatment and 8.98 for EPZ6438-treatment, which derives exactly from the jump in capture efficiency we discussed above. The good news for signal compression, however, is that it reduces the likelihood of mistakenly identifying off-target peaks as target as long as the user who paid for the data is willing to conclude there is no signal in the data. Thus, a data loss is incurred. It is our view that it would be better to see off-target peaks and know that they are off-target, as this constitutes useful information about the chromatin landscape. Before moving on, we note that the disparity between capture efficiency of the SNAP nucleosomes and cellular chromatin suggests that no region of the genome presents H3K27me3 consistently across all cells in the population. The SNAP spike-ins are homogeneous so they provide a great source of information on heterogeneity even though they have so far never been used in this way.

In Figure 2C, siQ-ChIP reveals a reduction of the right-most peak and an increase in the left-most peak for the region of chromosome 2 in the visualized window. The heat maps show clear density underlying both peaks, as well as the region between the peaks, in both experimental contexts. siQ-ChIP analysis can be used to diagnose the nature of these features. For example, Figure 6A-B show HMD and siQ-ChIP (*ê*(*x*)) on another region of chromosome 2, where each method reports a loss of signal for H3K27me3 after EPZ6438 treatment. HMD compresses the signal (post-EPZ6438) to such an extent that the level of background is different between treatment paradigms. Given that all data were collected the same way, using the same sequencer, it should be considered impossible to obtain drastically different background levels. The alternative is that the apparent background in DMSO-treated signal is an actual signal originated by antibody binding events, and that compression arises from off-target binding or a high degree of on-target heterogeneity post-EPZ6438. This is the ambiguous interpretation noted in the introductory remarks.

**Figure 6.**
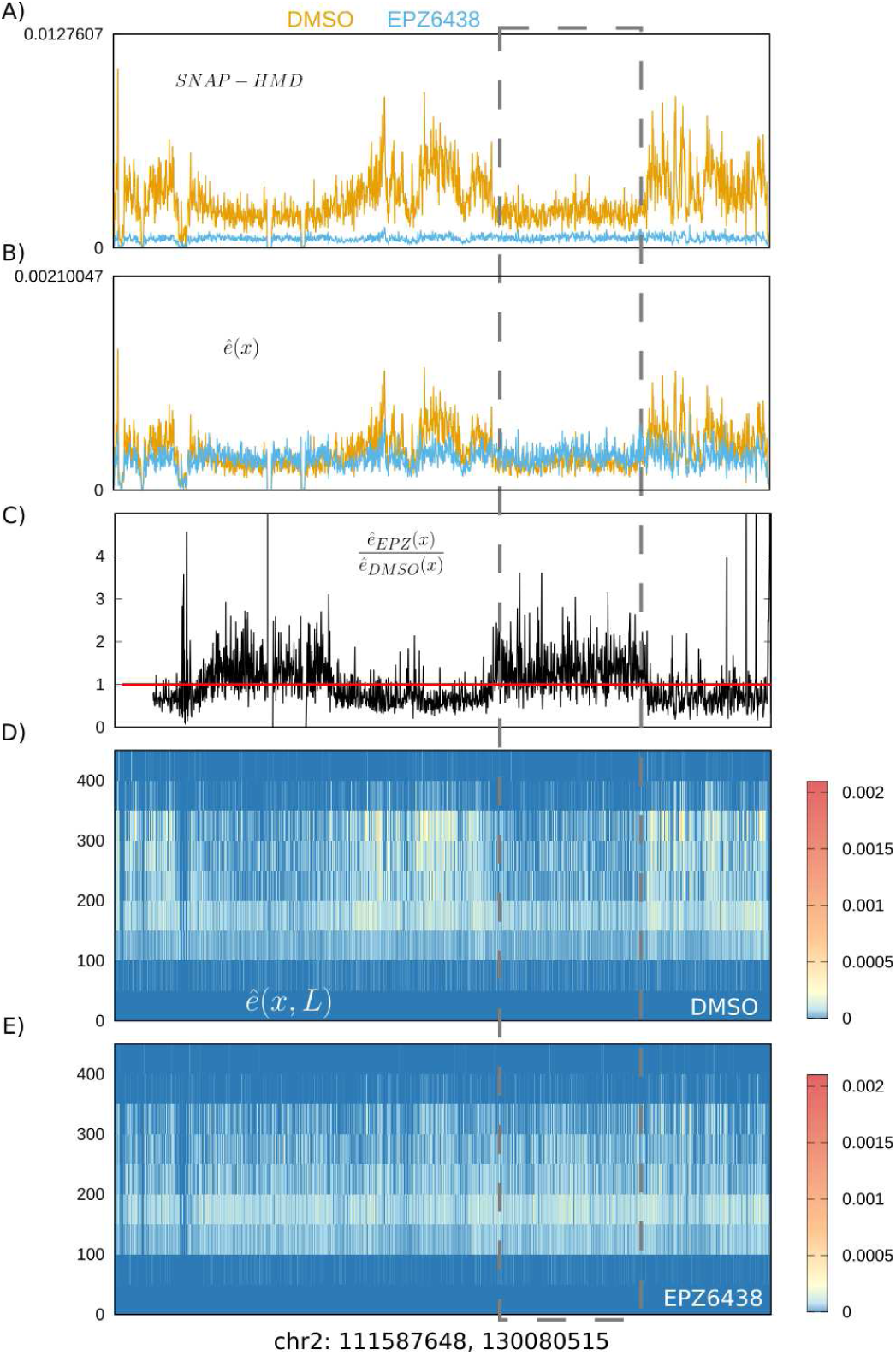
siQ-ChIP off-target detection. (A) HMD, (B) cumulative efficiency, (C) differential enrichment based on ratio of *ê*(*x*), and (D) *ê*(*x, L*). The red line is drawn at unity. Window width is 10k bp.

In Figure 6B, siQ-ChIP shows that peaks in DMSO-treatment are brought down to levels that might be consistent with background in both experimental conditions, which is what would be expected of real background (it should be similar in both data sets). However, siQ-ChIP provides a simple test of differential enrichment, shown in Figure 6C. This form of analysis is not possible with any other quantitation scheme. The ratio of cumulative efficiencies clearly shows both an EPZ6438-dependent loss in efficiency (regions lower than the red line) and an EPZ6438-dependent gain in efficiency (regions higher than the red line). Figure 6 also shows *ê*(*x, L*) to highlight the fact that in sequencing data for DMSO-treated samples, we observe non-zero capture of the genomic regions associated with enhanced efficiency post-EPZ6438 treatment. One such region is highlighted with a dashed box. Like HMD, but without signal compression, the genome-browser view of siQ-ChIP suggests that the enhanced capture post EPZ6438 treatment is due to either off-target binding or heterogeneous spread of H3K27me3. The siQ-ChIP object *ê*(*x, L*), which cannot be viewed in a browser, reveals something more about these regions of interest. We observe a reduction in warmer colors for dinucleosomes and an expansion of more dim shades of color exclusively for mononucleosomes. This implies a loss of tight binding and a gain in weak binding, that is not strongly enhanced by dinucleosomal avidity, because the efficiency is directly related to effective binding constants (Equation (12)). siQ-ChIP analysis is less vague in implication here, and suggests that these regions of differentially enhanced signal post EPZ6438 bear off-target PTMs, and that these off-targets may be found in DMSO signal as well.

The model above affords the following explanation for the behavior of capture efficiency: The concentration of free antibody is increased by epitope depletion, relative to the DMSO control, leaving the free antibody to bind off-target PTMs with higher frequency (but not better binding constants). This leads to the observed trends in differential enrichment where off-target regions are mildly enhanced after target depletion. If this implication can be proven, there are significant consequences. For example, if these regions turn out to derive from off-target binding then these off-target signals are often passing as background or are totally ignored upon global normalization, moreover the SNAP spike-ins have qualified this antibody as specific. With this in mind, we next sought to identify whether any antibody driven interactions might underlie our observations.

### Significant off-target enrichment in EPZ6438 treated chromatin

Meta analysis of existing ChIP-seq data in HCT116 colon cancer cells was used to investigate the potential for off-target signal in the above sequencing results. Correlation among sequencing peaks in different antibody tracks was evaluated. Overlap in coverage from multiple antibodies targeting different PTMs is taken as statistical evidence that the coverage could be due to either mark. Depending on the extent of overlap, as is shown below for H3K9me3, this is taken as strong evidence in support of off-target binding.

Figure 7 shows the two different regions of chromosome 2 that we have looked at in Figures 2 and 6. The siQ-ChIP differential enrichment is plotted along with tracks for H3K9me3 and H3K36me3. The H3K36me3 track is taken from ENCODE[12] (ENCSR091QXP, GEO:GSE95914), while the H3K9me3 track was generated in our lab, albeit for purposes unrelated to our present endeavor. It was basic human-level pattern recognition that initially clued us in on the overlap with H3K9me3. The presentation in Figure 7 clearly demonstrates a striking degree of overlap in regions enhanced by EPZ6438 treatment and H3K9me3, while showing that H3K36me3 is difficult to anticipate from the projected browser view. We know from *ê*(*x, L*) in Figure 2 that both DMSO and EPZ6438 treatments report weak capture that is consistent with H3K36me3, but it escapes easy detection when looking only at ratios of *ê*(*x*). Any off-target naturally overlapping with target will be difficult to detect with this simplistic approach. From Figure 5, one would expect H3K4me3, H3K36me3, and H4K20me3 to be among the most likely off-target epitopes, but would not have been considered a significant concern based on 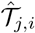.

**Figure 7.**
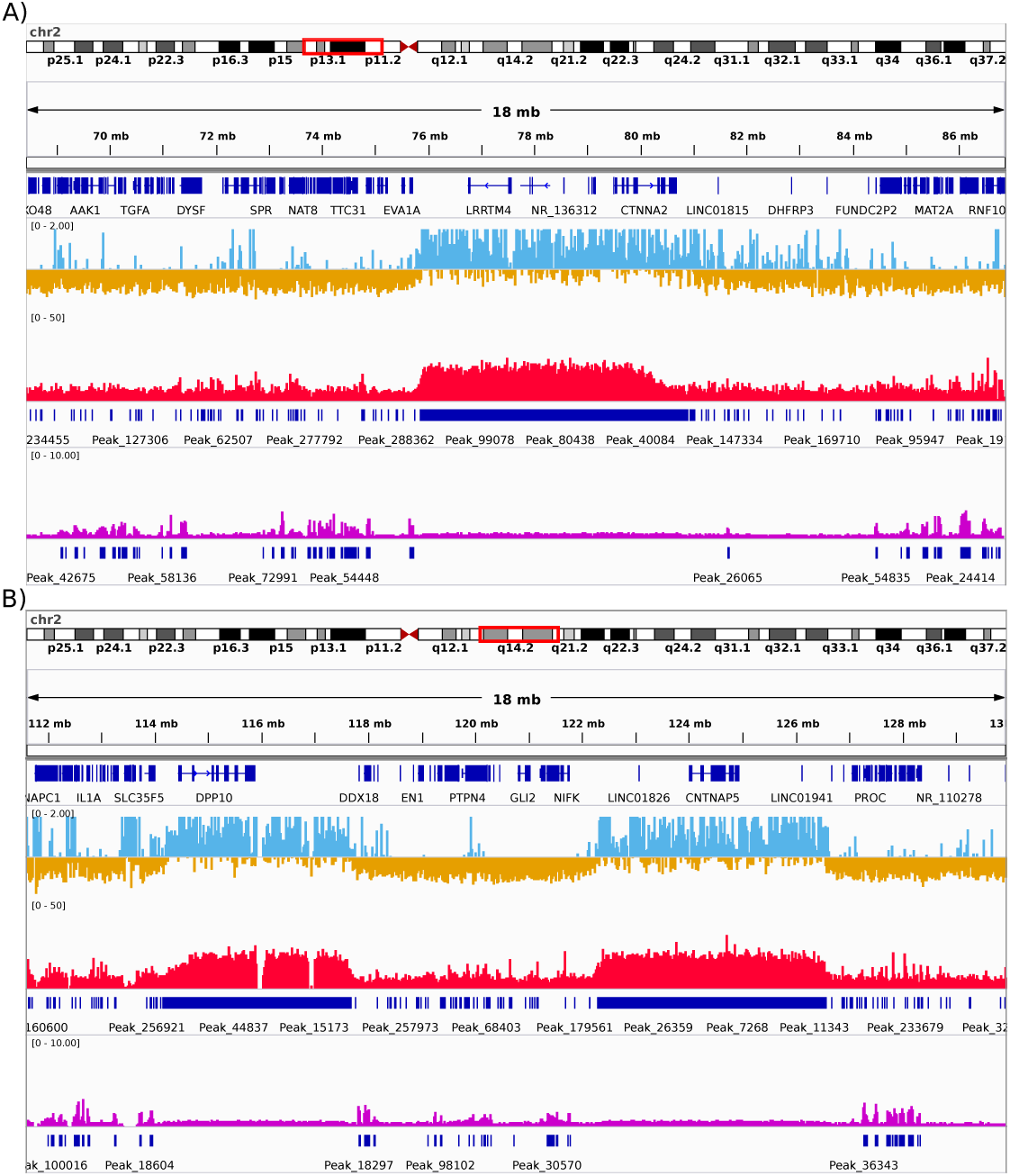
Browser views of siQ-ChIP enrichment and off-target tracks. Examples of K9me3 (red) and H3K36me3 (purple) tracks on chr2 in IGV genome browser. Cumulative efficiency *ê*(*x*) is computed on windows 2000 bases wide. Blue-Gold track is the ratio of EPZ6438 cumulative efficiency to DMSO cumulative efficiency. Blue color indicates higher efficiency in EPZ6438 treatment, Gold indicates higher efficiency in DMSO. The blue-gold track is siQ-ChIP differential enrichment. The red track is H3K9me3 ad the purple track is ENCODE H3K36me3. A) The region of chr2 shown in figure 2. B) The region of chr2 shown in figure 6.

To quantify the observed overlap globally, we recorded the total genomic coverage of peaks in either H3K9me3 or H3K36me3 ChIP-seq experiments and asked how frequently DMSO or EPZ6438 ChIP-seq peaks covered the same regions. To avoid dependence on siQ-ChIP while establishing the extent of overlap, we simply called peaks in each dataset using MACS2[17]. Given our analysis above, we expected a number of off-target binding events to be classified as background, so we suggest this peak-coverage analysis is a lower bound for what is actually present in the data. Tables 2 and 3 summarize our results.

**Table 2.** Peak numbers and Hyper-geometric P-values (left) for genome wide signal overlaps. 0 indicates significant under sampling, 1 indicates significant over sampling.

|  | K9me3 | K36me3 | DMSO<sub>MACS</sub> | EPZ6438<sub>MACS</sub> |
| --- | --- | --- | --- | --- |
| K9me3 | – | 0 | 1 | 1 |
| K36me3 | 0 | – | 1 | 1 |
| DMSO<sub>MACS</sub> | 1 | 1 | – | 1 |
| EPZ6438<sub>MACS</sub> | 1 | 1 | 1 | – |

**Table 3.** Enrichment bias as fold change for genome wide signal overlaps

|  | K9me3 | K36me3 | DMSO<sub>MACS</sub> | EPZ6438<sub>MACS</sub> |
| --- | --- | --- | --- | --- |
| K9me3 | – | 0.65 | 1.33 | 4.19 |
| K36me3 | 0.65 | – | 1.92 | 2.77 |
| DMSO<sub>MACS</sub> | 1.33 | 1.92 | – | 6.13 |
| EPZ6438<sub>MACS</sub> | 4.19 | 2.77 | 6.13 | – |

Table 3 reports that, for MACS-called peaks, H3K9me3 overlaps with H3K36me3 coverage 0.65 times less than would be expected by chance. DMSO-treated chromatin that was immunoprecipitated with the H3K27me3 antibody overlaps with H3K9me3 1.33 times more often than expected when peaks are called by MACS. IP with the H3K27me3 antibody produced significant overlap with H3K36me3 coverage for DMSO-treated cells.

The amount of overlap with off-target H3K9me3 or H3K36me3 shows a substantial increase in the EPZ6438-treated chromatin relative to DMSO. This reinforces the model prediction that, even though the antibody is specific, it will IP only off-target species if the target epitope is removed. Alarmingly, 43% of the MACS-called IP coverage in EPZ6438 ChIP-seq are correlated to either H3K9me3 or H3K36me3, and two of the strongest off-targets (by Figure 5, H3K4me3 and H4K20me3) are yet to be compared to the H3K27me3 track. These observations strongly imply that the peaks revealed by ChIP-seq of EPZ6438 treated cells cannot be interpreted as peaks of migrated or maintained H3K27me3.

The overlap data also shows a significant correlation between DMSO and EPZ6438 data based on MACS peak calling. Interestingly, 47.5% of this overlap also overlaps with H3K9me3 coverage and an additional 5% overlaps with H3K36me3 coverage — Over half of the DMSO and EPZ6438 coverage overlap is correlated with off-target PTM coverage. These overlapping regions are therefore ambiguously determined as either H3K27me3, H3K9me3, or H3K36me3 bearing regions. Overall, the peaks in EPZ6438 data are now much more easily interpreted as off-target than on-target.

### Flexible protocols

For hard to ChIP PTMs it may be impossible to obtain a workable amount of DNA from one IP reaction. It is possible to combine *M* IP reactions, where each IP is required to have identical reaction conditions, as follows

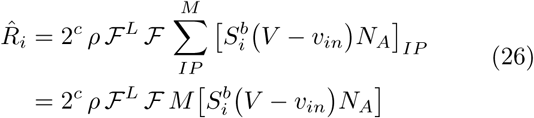

Using this adjustment in the effective efficiency only modifies Equation (23) by the factor 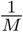. This result is intuitive, as it clearly maps the results on to a per-IP basis. Explicitly, the effective efficiency becomes

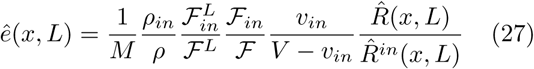

Obviously, normalization to cell number is encoded here as well.

## Conclusion

We have developed a completely spike-in free quantitative analysis for ChIP-seq. The approach acknowledges that the IP is a binding reaction and employs the common framework for binding reactions to yield quantitative relationships for sequencing results. We have shown that projecting to the standard browser-style view collapses the three dimensional map of capture efficiency *ê*(*x, L*) into a less informative representation, and that *ê*(*x, L*) is a useful object for suggesting whether a browser peak is due to target or off-target binding. siQ-ChIP applies to standard paired-end MNase or crosslinking ChIP protocols and only requires that each step of the process be carefully logged so that the scale can be correctly determined. MNase data is likely to produce more powerful insights as it allows one to distinguish mono- and di-nucleosome binding events.

We have used the framework of siQ-ChIP to investigate SNAP-HMD and ChIP-based assessment of antibody specificity. Predictions were tested for both capture efficiency and specificity 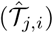, showing that our model is fully capable of describing chromatin spikeins. The behavior of these spike-ins is not fully appreciated in the literature, and our analysis places new limitations on the utility of “specificity” measures. In particular, we revealed significant off-target binding for an antibody that displays a high degree of specificity within the SNAP-HMD metrics. We also suggest that differences in heterogeneity between the spike-in and cellular chromatin of interest cause an unappreciated compression of signal.

Our work may also hold implications for state-of-the-art peak callers that infer/learn background levels by analyzing signal. We have shown that what seems like background in DMSO signal displays a response to epitope depletion and correlates significantly with H3K9me3 domains. Thus, this “background” is actually likely to be residual off-target capture, even in DMSO-derived data.

The above work also suggests that any ChIP-seq data should be cross-validated against a mixture of tracks from the same cell line to help ensure a lack of ambiguity in any peaks that are designated to a particular PTM. As the possibility of PTM co-occurence is valid, we argue that co-designation of peaks is more appropriate than mis-designation. In the case presented above, the overlap with H3K9me3 is striking, and we suggest that co-occurance everywhere must have smaller odds than cross-reactions.

We have also noted some limitations in the siQ-ChIP differenential enrichment analysis, where any off-target that is in two samples may be difficult to see. Future work will investigate using hidden Markov models to find these sorts of features within the differential enrichment quotient. The above work also opens a number of doors toward optimizing the information per dollar per lab hour spent on ChIP-seq.

Finally, we close with a proposition for profiling antibodies. Equation (15) can be used to empirically determine relative binding constants for different epitopes if an experiment can be designed such that the observable fraction of each epitope is equal. Equality in observability can be achieved by replacing cellular chromatin with unmodified synthetic nucleosomes. The PTM bearing spike-in nucleosomes would then have equal observabilities. A standard ChIP-seq with the antibody of choice can then be analyzed using Equation (15) where *e*_*i*_ is given by Equation (18). This approach would provide a profile of the antibody binding constants rather than the contextual profiles shown in figure 4.

The software to compute the siQ-ChIP prefactors and objects *ê*(*x*) and *ê*(*x, L*) is available at GitHub[18]. Documentation is also found there.

## Author’s contributions

B.M.D. developed theory, software, designed and analyzed experiments, and wrote the manuscript. R.L.T. performed experiments and sequencing analysis and contributed many practical insights. R.M.V. and A.A.C. performed experiments. E.M.C. performed numerous pilot experiments. R.M.V. and E.M.C. also contributed to many discussions. S.B.R. designed experiments and wrote the manuscript.

## Acknowledgements

We thank Tim Triche Jr., and Benjamin K. Johnson for helpful comments and discussions. This work was supported in part by grants R35GM124736 (S.B.R.) and F32CA225043 (A.A.C.) from the National Institutes of Health, and grant PF-16-245-01-DMC (R.L.T.) from the American Cancer Society - Michigan Cancer Research Fund.

## Tables

**Additional Files**

Additional file 1 — Experimental methods:

PDF file containing details of experimental protocols. Code and scripts are published at GitHub[18]. GEO accession number is forthcoming.

## Supplemental Methods and Figures

**Table S1.** Required measurements for running siQ-ChIP algorithm

**Table S2.** H3K27me3-positive and negative regions

**Figure S1.** H3K27me3 Native ChIP optimizations

**Figure S2.** Antibody-chromatin complex formation binding reaction

### Native Chromatin Immunoprecipitation (ChIP)

HCT116 colorectal carcinoma cells were treated for 72 hours with either vehicle (0.02% DMSO and 0.03% PBS) or EZH2 inhibitor (EPZ6438, 1 uM). Cells were trypsinized, collected, flash frozen, and stored at −80°C.

#### Nuclei purification

To purify nuclei, thawed cells were washed 3x with PBS, 2x with Buffer N (15 mM Trizma Base pH 7.5, 15 mM NaCl, 60 mM KCl, 5 mM MgCl2, 1 mM CaCl2, 8.5% sucrose, 1 mM DTT, 200 uM PMSF, 50 ug/mL BSA, and protease inhibitor), and lysed with 2x Lysis Buffer (0.6% NP-40 in Buffer N). Nuclei were layered over a 30% sucrose cushion, pelleted at 1300 rcf, and resuspended in Buffer N.

#### Nucleosome preparation

Crude chromatin concentration was determined by sonicating 2 uL of nuclei in 18 uL NaCl (2 M) and measuring DNA concentration with a Nanodrop; nuclei equivalent to 50 ug of DNA were aliquoted and spiked with 5 uL of a SNAP-ChIP k-MetStat panel (Epicypher). 1 U MNase (25 U/uL) was added per 4.275 ug chromatin and incubated at 37°C for 12 minutes with shaking. MNase digestion was stopped with 1/10 volume MNase Stop Buffer (10x, 0.1 M EGTA) followed by 1/8 volume NaCl (5 M) to lyse nuclei and release digested chromatin. After centrifugation, soluble chromatin was added to 33 mg of rehydrated ceramic hydroxyapatite (CHT) resin and rotated for 10 min at 4°C. The CHT Resin:Chromatin mix was added to a centrifugal filter unit and washed 4x with HAP Wash Buffer 1 (5 mM NaPO4 pH 7.2, 600 mM NaCl, 1 mM EDTA, and 200 uM PMSF), 4x with Hap Wash Buffer 2 (5 mM NaPO4 pH 7.2, 100 mM NaCl, 1 mM EDTA, and 200 uM PMSF), and eluted with three successive additions of HAP Elution Buffer (500 mM NaPO4 pH7.2, 100 mM NaCl, 1 mM EDTA, and 200 uM PMSF). DNA concentration was again measured by sonication and adjusted to 20 ug/mL with ChIP Buffer 1 (25 mM Tris pH 7.5, 5 mM MgCl2, 100 mM KCl, 10% glycerol, 0.1% NP-40, 200 uM PMSF and 50 ug/mL BSA).

#### Antibody:bead preparation

Antibody:bead conjugates were prepped by adding 5 or 10 uL of Cell Signaling Technology H3K27me3 antibody (CST #9733, clone C36B11, lot 14, 102 ug/mL) or 3 uL of abcam H3K27me3 antibody (ab6002, 1 mg/mL) to 12.5 uL of Protein A Magnetic Dynabeads (Invitrogen) that had been washed with ChIP Buffer 1. 5 uL of CST #9733 was chosen as the optimal antibody after comparison with ab6002 (Fig. S1A). Antibody:bead conjugates were incubated on a rotator for 3 hours at 4°C, washed 2x, and resuspended with ChIP Buffer 1.

#### Chromatin Immunoprecipitation

Between 0.5 and 3 ug of chromatin was used for optimization (Fig. S1B). Based on this, 0.75 ug of the purified digested chromatin (as measured by DNA concentration) was added to the antibody:bead conjugates while a volume equivalent to 10% was saved for input from the purified chromatin. The bead:antibody:chromatin mixture volume was brought to 100 uL with ChIP Buffer 1 and incubated on a rotator for 17 min at 4°C. Using a magnetic rack, the mixture was washed for 10 min on a rotator at 4°C 2x with ChIP Buffer 2 (25 mM Tris pH 7.5, 5 mM MgCl2, 300 mM KCl, 10% glycerol, 0.1% NP-40, 200 uM PMSF and 50 ug/mL BSA), 1x with ChIP Buffer 3 (10 mM Tris pH 7.5, 250 mM LiCl, 1 mM eDTA, 0.5% NaDeoxycholate, 0.5% NP-40, 200 uM PMSF and 50 ug/mL BSA), 1x with ChIP Buffer 1, 1x with TE Buffer (pH 8.0), and resuspended in 50 ul of ChIP Elution Buffer. This was incubated at 55°C for 5 min, and sample was eluted from beads on a magnetic rack. Finally, 2 uL NaCl (5M), 1 uL EDTA (0.5M), and 0.5 uL Proteinase K (100x) were added to ChIP and Input samples, which were incubated overnight at 55°C.

#### Purification of Immunoprecipitated DNA

DNA was recovered using KAPA Pure beads at a 1.5X ratio on a magnetic rack. After two washes with 75% EtOH, DNA was eluted in 50 uL dH20 and DNA concentration was measured with a Qubit. Input and ChIP Libraries were prepared from 10 ng DNA using a KAPA Hyper Prep Kit. Libraries were purified once more with KAPA Pure beads at a 1.0X ratio to remove adaptor contamination.

See Table S1 for measurements recorded during ChIP experiments that are used to calculate the estimated capture efficiency with the siQ-ChIP algorithm.

**Table S1.** Required measurements for running siQ-ChIP algorithm

| *Symbol* | *Measurement* | *Veh1* | *Veh2* | *EPZ1* | *EPZ2* | *Sample Equation (e.g. Veh1)* |
| --- | --- | --- | --- | --- | --- | --- |
| $\mathcal{F}$ | Total IP mass (ng) | 55.50 | 56.50 | 24.20 | 35.20 | =10/55.5 |
|  | IP mass into library (ng) | 10 | 10 | 10 | 10 |  |
| $\mathcal{F}_{in}$ | Total input mass (ng) | 57.50 | 57.50 | 50.50 | 50.50 | =10/57.5 |
|  | Input mass into library (ng) | 10 | 10 | 10 | 10 |  |
| $\mathcal{F}^L$ | *IP library conc. (nM) | 55.75 | 56.60 | 42.80 | 59.26 | =(10*2)/(55.75*20) |
|  | Total IP library volume (ul) | 20 | 20 | 20 | 20 |  |
|  | *Normalized IP library conc. (nM) | 10 | 10 | 10 | 10 |  |
|  | *Normalized IP volume (ul) | 2 | 2 | 2 | 2 |  |
| $\mathcal{F}_{in}^L$ | *Input library conc. (nM) | 121.76 | 121.76 | 85.15 | 85.15 | =(10*4)/(121.76*20) |
|  | Total input library volume (ul) | 20 | 20 | 20 | 20 |  |
|  | *Normalized input library conc. (nM) | 10 | 10 | 10 | 10 |  |
|  | *Normalized input volume (ul) | 4 | 4 | 4 | 4 |  |
| $V_r$ | Volume taken for input (*vin*) (ul) | 100 | 100 | 100 | 100 | =(100)/(1000-100) |
|  | Total volume of chromatin (*V*) (ul) | 1000 | 1000 | 1000 | 1000 |  |
| $\rho$ | Mass into library (ng) | 10 | 10 | 10 | 10 | **expectedIP**=(1e-09*10*(2<sup>11</sup>))<br>/(691*660<sup>#</sup>)/(20*1e-6)<br>=1/(expectedIP/55.75) |
|  | PCR amplifications (Ct) | 11 | 11 | 11 | 11 |  |
|  | Total library volume (ul) | 20 | 20 | 20 | 20 |  |
|  | *IP library concentration (nM) | 55.75 | 56.60 | 42.80 | 59.26 |  |
|  | *Average IP fragment length (bp) | 691 | 735 | 648 | 676 |  |
| $\rho_{in}$ | Mass into library (ng) | 10 | 10 | 10 | 10 | **expectedIN**=(1e-09*10*(2<sup>11</sup>))<br>/(367*660<sup>#</sup>)/(20*1e-6)<br>=1/(expectedIN/121.76) |
|  | PCR amplifications (Ct) | 11 | 11 | 11 | 11 |  |
|  | Total library volume (ul) | 20 | 20 | 20 | 20 |  |
|  | *Input library concentration (nM) | 121.76 | 121.76 | 85.15 | 85.15 |  |
|  | *Average input fragment length (bp) | 367 | 367 | 380 | 380 |  |

\*atypical ChIP measurement to be collected by experimentalist
<sup>#</sup>660 g/mol is the average molecular weight per base pair of DNA

### q-RTPCR

To optimize the ChIP experiment, positive genomic regions reportedly marked by H3K27me3 (HCT116 ENCODE data; GEO: GSE86755) and negative regions lacking H3K27me3 were tested by q-RTPCR using SYBR mastermix and forward and reverse primers for each region (Table S2).

**Table S2.**
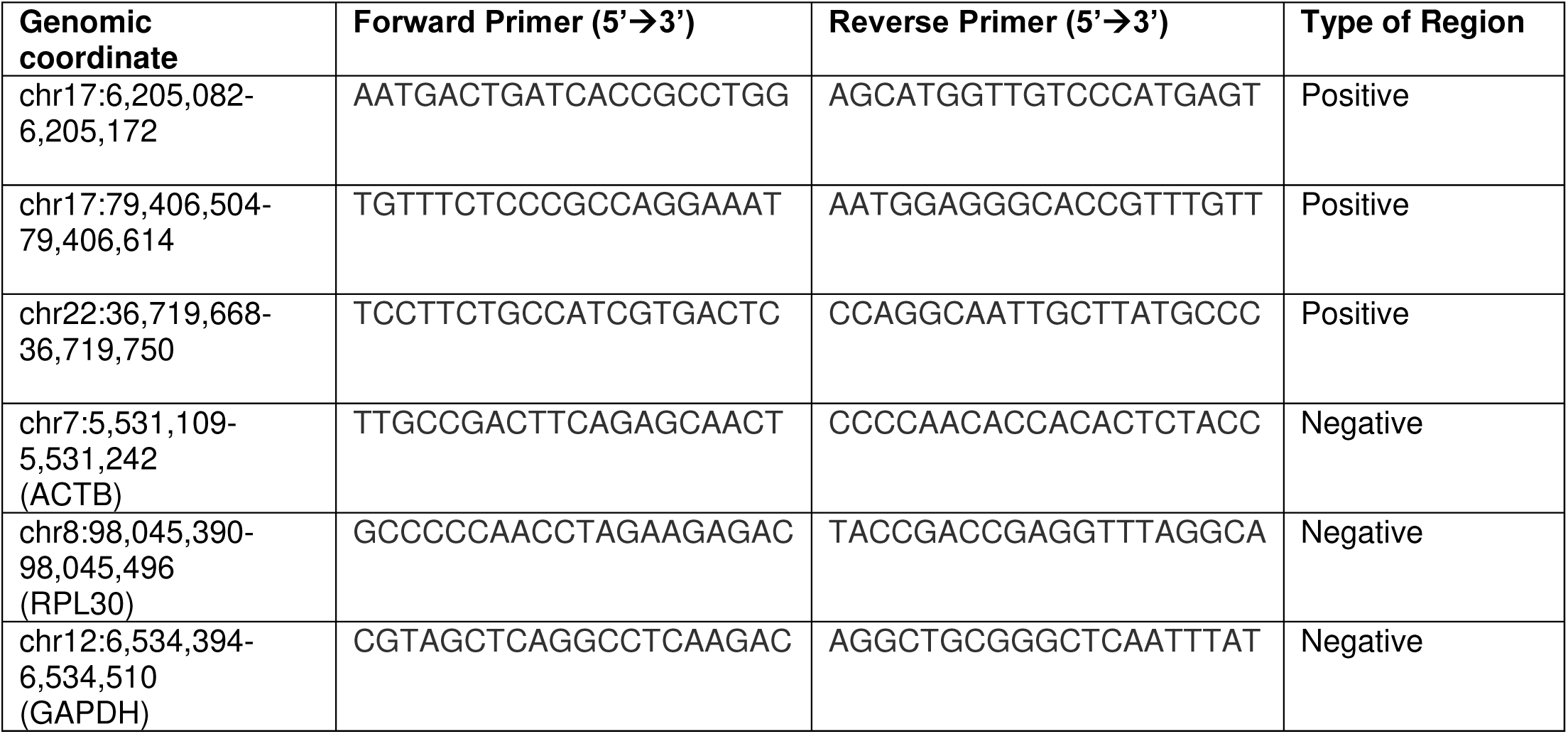
Primers for H3K27me3 positive and negative regions

### Sequencing

Prepared libraries were submitted to the VARI Genomics Core for library QC and quantification using the Agilent 2100 Bioanalyzer and KAPA Library Quantification Kit, respectively. Libraries were sequenced on an Illumina NextSeq 500 with 2 × 75 bp paired-end reads.

### NGS Data Preprocessing

Adapters were trimmed from fastq sequences using TrimGalore! version 0.5.0 (https://github.com/FelixKrueger/TrimGalore). Trimmed sequences were queried for overall sequencing quality using FastQC version v.0.11.8 (https://www.bioinformatics.babraham.ac.uk/projects/fastqc/). Sequences were then aligned to the human hg38 genome build using the following command in bowtie2 version 2.3.4.3 (PMID: 22388286):

bowtie2 -I 0 -X 700 --end-to-end --sensitive -x /path_to_hg38_index -1

/path_to_first_paired_read.fastq.gz -2 /path_to_second_paired_read.fastq.gz -S sample.sam

SAM files were further processed with the following command to isolate paired reads with high mapping quality, correct pair orientation, and to calculate fragment length:

awk -v MAQ=20 ‘$5>=MAQ && $2==99 ‖ $5>=MAQ && $2==163 {print $3”\t”$4”\t”$4+$9-1} sample.sam | awk ‘$2<=$3{print $1”\t”$2”\t”$3”\t”$3-$2} | sort -k1,1 -k2,2n > outfile.bed

Finally, known blacklisted regions were removed from the bed files using the subtract function from bedtools (PMID: 24782889, 20110278).

#### MACS2 peak calling

To call peaks of modification enrichment for both our H3K9me3 ChIP-seq and H3K27me3 ChIP-seq datasets, we used the following command from macs2 (v2.1.2):

macs2 callpeak -t {IP_sample}.bam -c {Input_sample}.bam -f BAMPE -n {SAMPLE}–outdir /path_to_output_dir/--broad -B

#### Bedtools intersection

To determine the coverage of the genome (in bp) that intersected between two datasets we performed the following command in bedtools for the different datasets:

bedtools intersect -a dataset1.bed -b dataset2.bed -wo | awk’{sum += $column_with_overlap_bp_count;} END {print sum;}’ “$@”

### Chromatin-antibody Binding Assay

Antibody+chromatin interaction was measured (Fig. S2) on a MicroCal PEAQ-ITC (Malvern) at 4°C to investigate relevance of equilibrium models. HeLa poly-nucleosomes (EpiCypher, 16-0003) or H3K27me3 antibody (CST #9733) were diluted in ChIP Buffer 1 to 0.75 ug in 280 uL (cell volume) or 5 uL into 40 uL (syringe), respectively. After an initial delay of 150 seconds, a single 40 uL injection was performed over 8 seconds, followed by an equilibration period. Equilibrium is reached in approximately three minutes, showing that IP conditions are compatible with equilibrium binding models.

**Figure S1.**
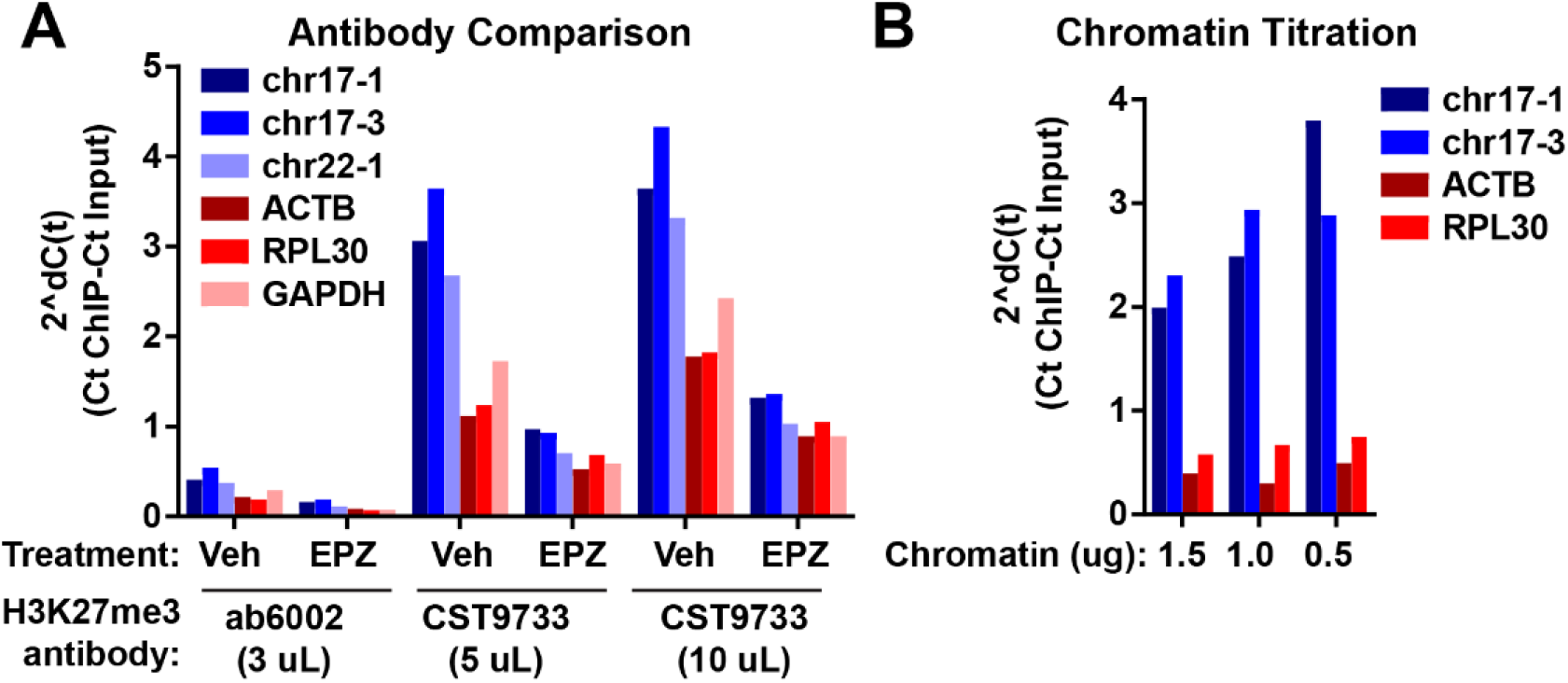
ChIP Optimizations. A) ChIPs from vehicle and EPZ6438 (EPZ) treated samples (chromatin equivalent to 3 ug DNA) using Abcam (ab6002) and Cell Signaling Technologies (CST #9733) H3K27me3 antibodies were compared. H3K27me3-positive (blue: chr17-1, chr17-2, chr22-1) and negative regions (red: ACTB, RPL30, and GAPDH promoters) were tested by qPCR. CST#9733 pulled down H3K27me3-positive regions more robustly. B) Different concentrations of chromatin equivalent to 1.5, 1.0, and 0.5 ug DNA were immunoprecipitated with CST #9733 (5 uL). Average of technical duplicates plotted for each.

**Figure S2.**
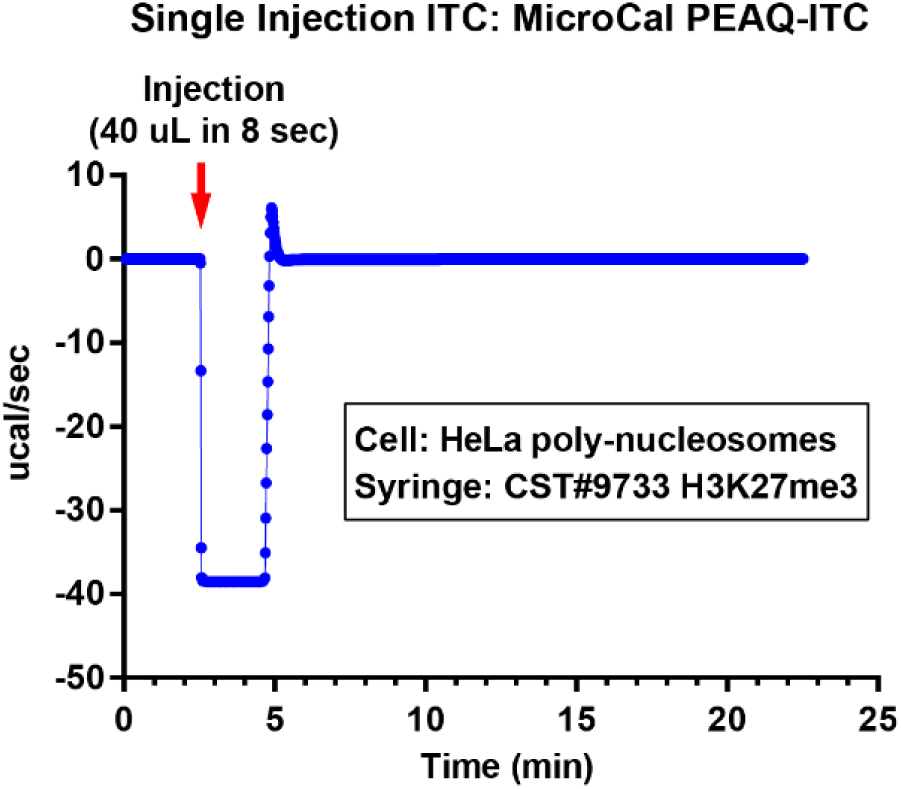
To evaluate whether the antibody-chromatin complex formation behaves as an equilibrium binding reaction, we performed a single injection isothermal titration calorimetry (ITC) experiment under the same conditions as a standard ChIP. H3K27me3 antibody and HeLa chromatin were each diluted in ChIP Buffer 1 and completely combined with a single injection on a microcalorimeter. After approximately three minutes, the binding reaction reached equilibrium.

## Footnotes

[1] If the efficiency ϵ is a known function of sequence *x*, then amplification can be more accurately accounted for with 1 + ϵ(*x*). Since ϵ is not usually known, we take the typical assumption 1 + ϵ(*x*) ≈ 2.

[2] The average base pair has molar mass 660 g/mole. 300 base pair fragments are then expected at 198000 g/mole. If 10 ng of 300 base pair long DNA are amplified by 2^11^ and suspended in 20 *µ*L (our library volume), the expected concentration is 5.17 *µ*M. *ρ* is the ratio of actual library concentration to the estimated 5.17 *µ*M. In practice, the empirical library averaged fragment length must be used to determine *ρ*. Thus, average fragment length is an input parameter for the software associated with this work.

